## Supplementary Information for "Global determinants of insect mitochondrial genetic diversity"

French et al.

##### Supplementary Fig. 1

The observed distribution of GDM (a, c, e, g, i, k) and GDE (b, d, f, h, l) for datasets with a minimum OTU per-cell threshold of 10 (a, b), 25 (c, d), 50 (e, f), 100 (g, h), 150 (i, j), and 200 (k, l). The number of sampled cells is 957, 572, 376, 245, 189, and 160, respectively for each OTU threshold.

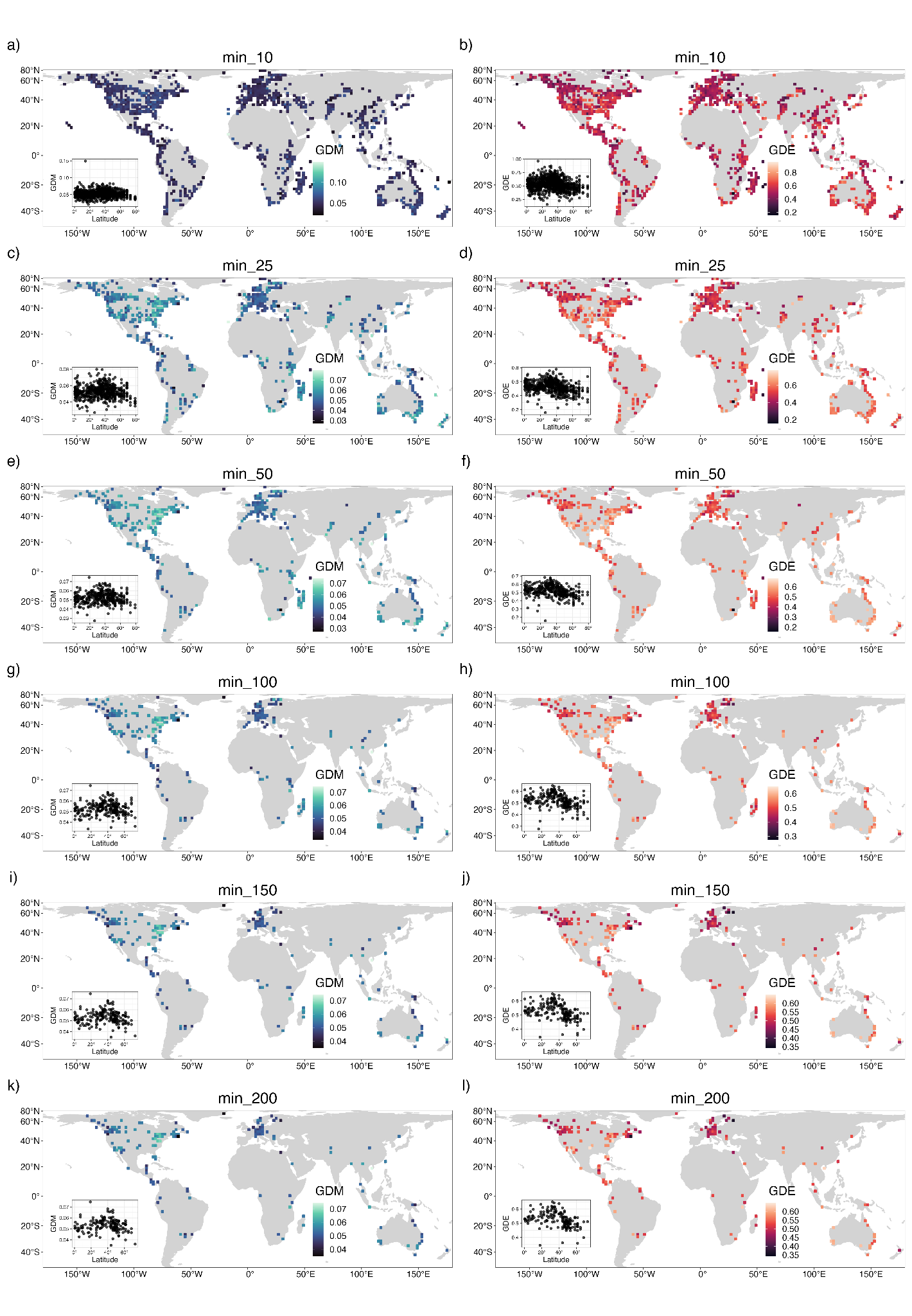

##### Supplementary Fig. 2

Criteria used to select the grid cell resolution and minimum number of OTUs per cell for this study. The three resolutions considered were 96.5 km x 96.5 km (“high”), 193 km x 193 km (“medium”), and 385.9 km x 385.9 km (“low”) equal-area grid cells using a Behrmann cylindrical equal-area projection. Each plot has the spatial resolution along the x-axis and the total number of cells along the y-axis. We additionally screened for a sampling scheme that balances the number of grid cells (a), median number of orders (b), variation in the number of OTUs across cells (c), median number of orders per cell (d), and total number of OTUs (e).
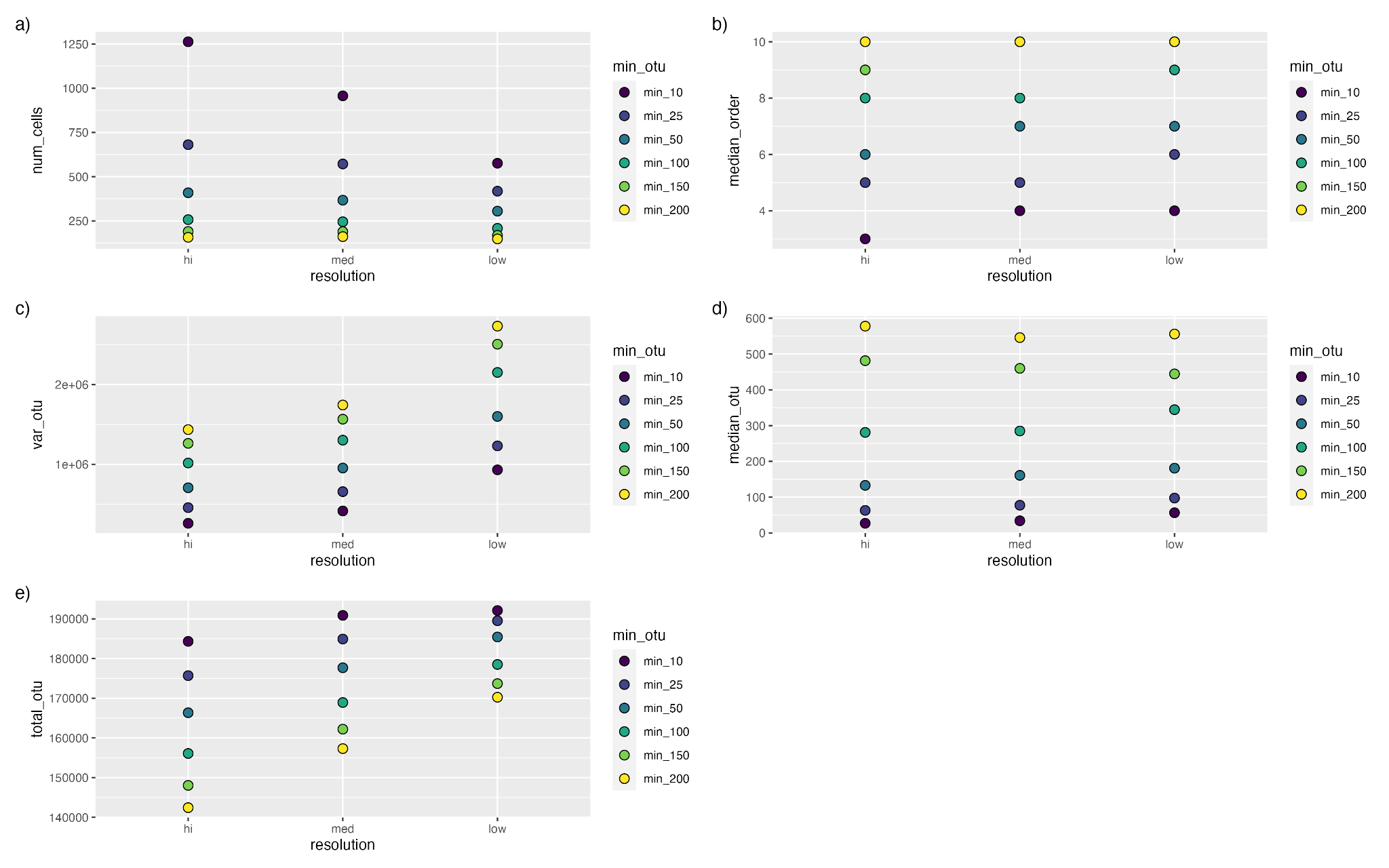

##### Supplementary Fig. 3

Observed data versus predicted data from GLMM models trained on 75% of data with 25% of the data withheld. Black lines indicate a perfect 1-to-1 relationship between observed and predicted data, and red lines with gray shading are trend lines with 95% confidence intervals from a regression between observed and predicted data. Points are colored according to the continent they were sampled in, demonstrating little spatial bias in residuals. Below each plot are summary statistics indicating model fit, where RMSE is the root mean squared error and Y-int is the y-intercept. Lower RMSE and higher R^2^ values indicate strong explanatory power, while a slope near one and a y-intercept near zero indicate low prediction bias.

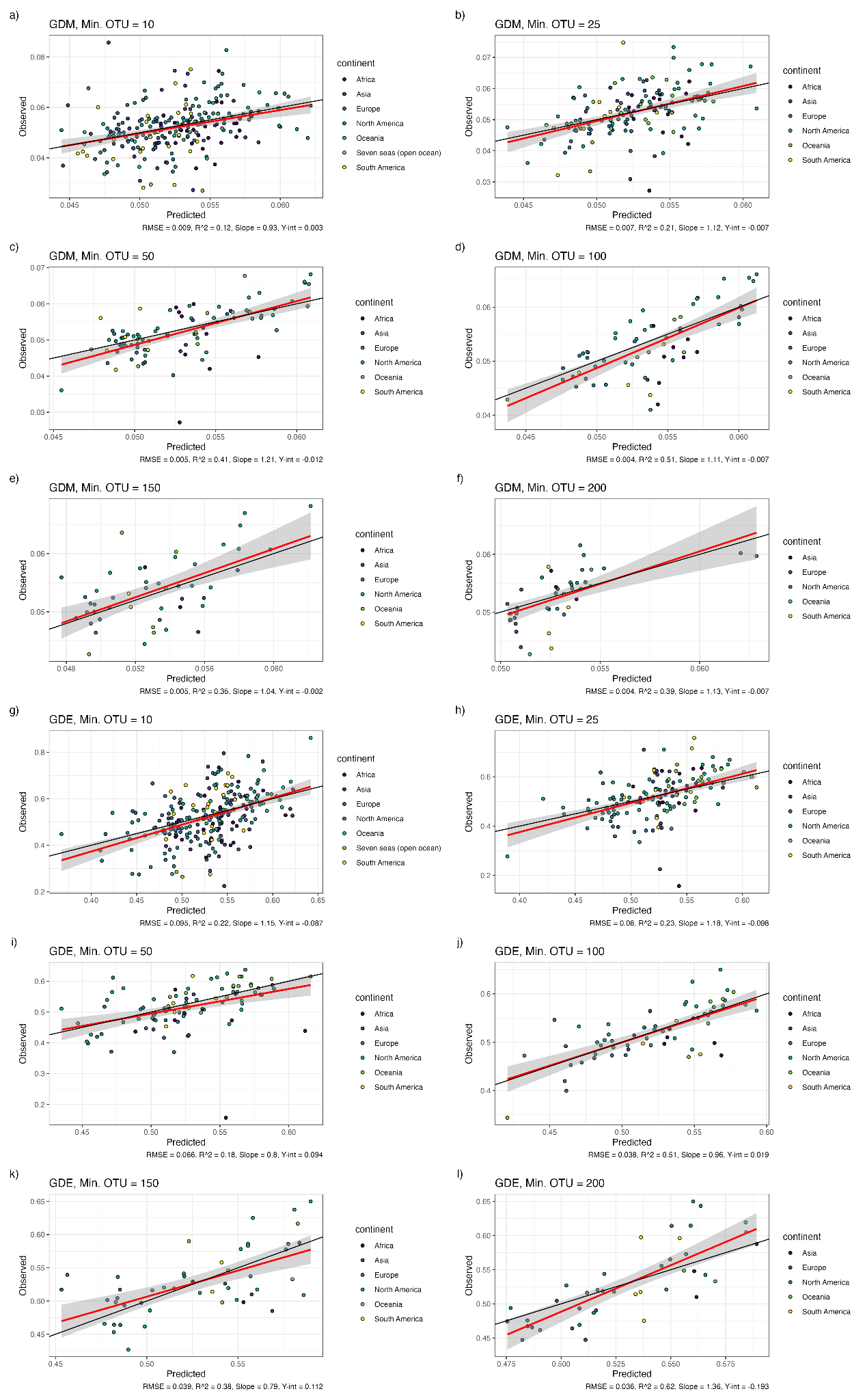

##### Supplementary Fig. 4

Global hotspots and coldspots of genetic diversity. These maps indicate grid cells in the top (red) and bottom (blue) 10% quantile of GDM and together (a-d), GDM alone (e-h), and GDE alone (i-l). Panels on the left are projections of the summary statistics across the globe from a spatial Bayesian GLMM model using environmental predictors (see Results) and panels on the right depict observed data. In the left-hand panels, areas with environments non-analogous to the environments used for modeling were removed prior to calculating the 10% quantiles.

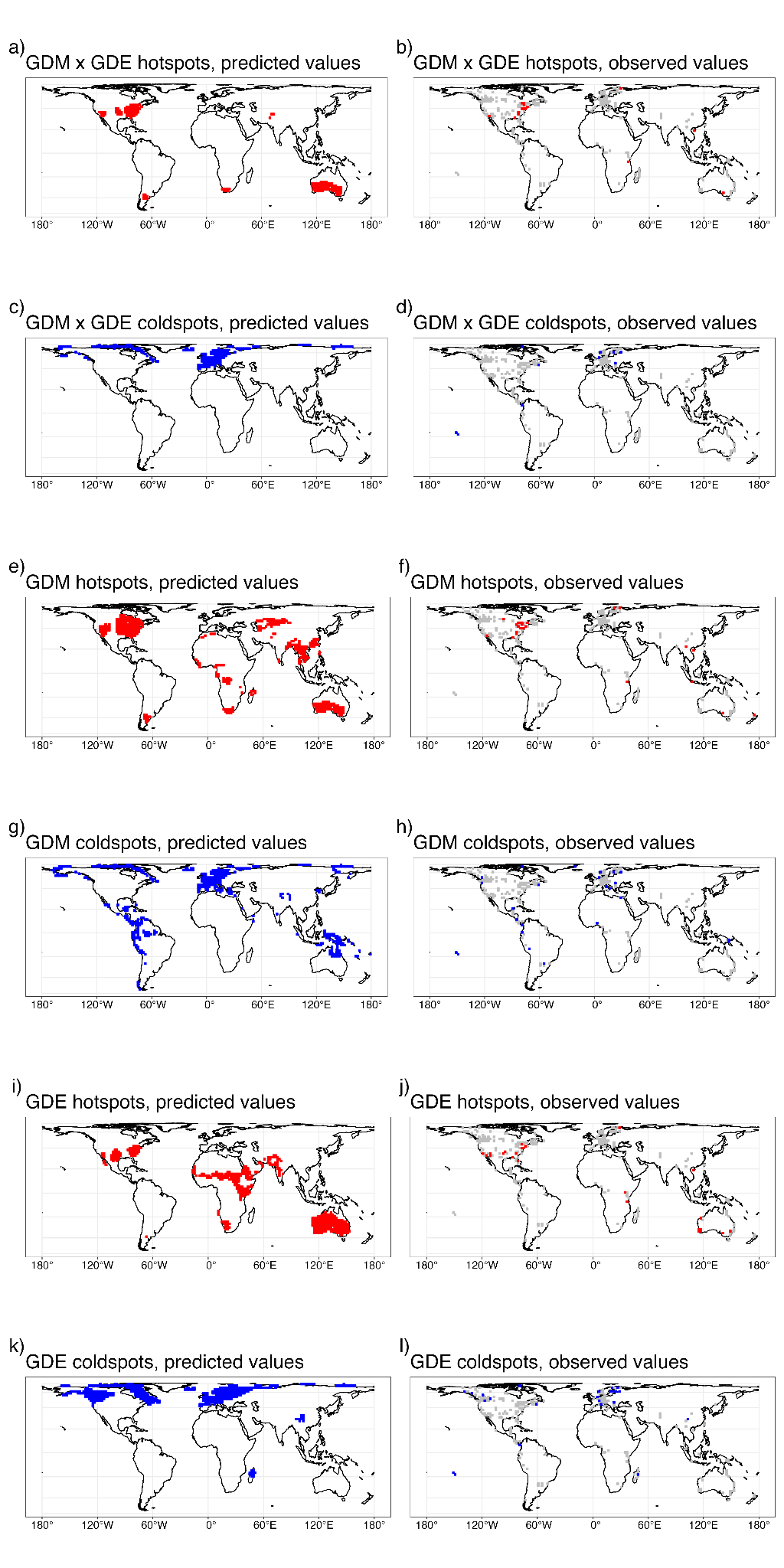

##### Supplementary Fig. 5

Global maps of the top environmental predictors of GDM and GDE. MTWM is the maximum temperature of the warmest month bioclimatic variable (°C * 10) (a), Temp. Trend describes centennial trends in temperature variation since the last glacial maximum (LGM) (b), PWM is the precipitation of the wettest month bioclimatic variable (mm) (c), Precip. Trend describes centennial trends in precipitation variation since the LGM (d), Temp. Range is the mean diurnal air temperature range bioclimatic variable (ºC * 10) (e), Precip. Seasonality is the precipitation seasonality bioclimatic variable (kg m^-2^) (f), Precip. Variation is the standard deviation around centennial trends in precipitation variation since the LGM (g), and PDM is the precipitation of the driest month bioclimatic variation (kg m^-2^) (h). Precip. Seasonality, PWM, PDM, Temp. Trend, Precip. Variation, MTWM, Temp. Range, and Precip. Trend were the eight predictors in a Bayesian GLMM model of GDM, while Temp. Trend, PWM, and MTWM were the three predictors in a model of GDM.

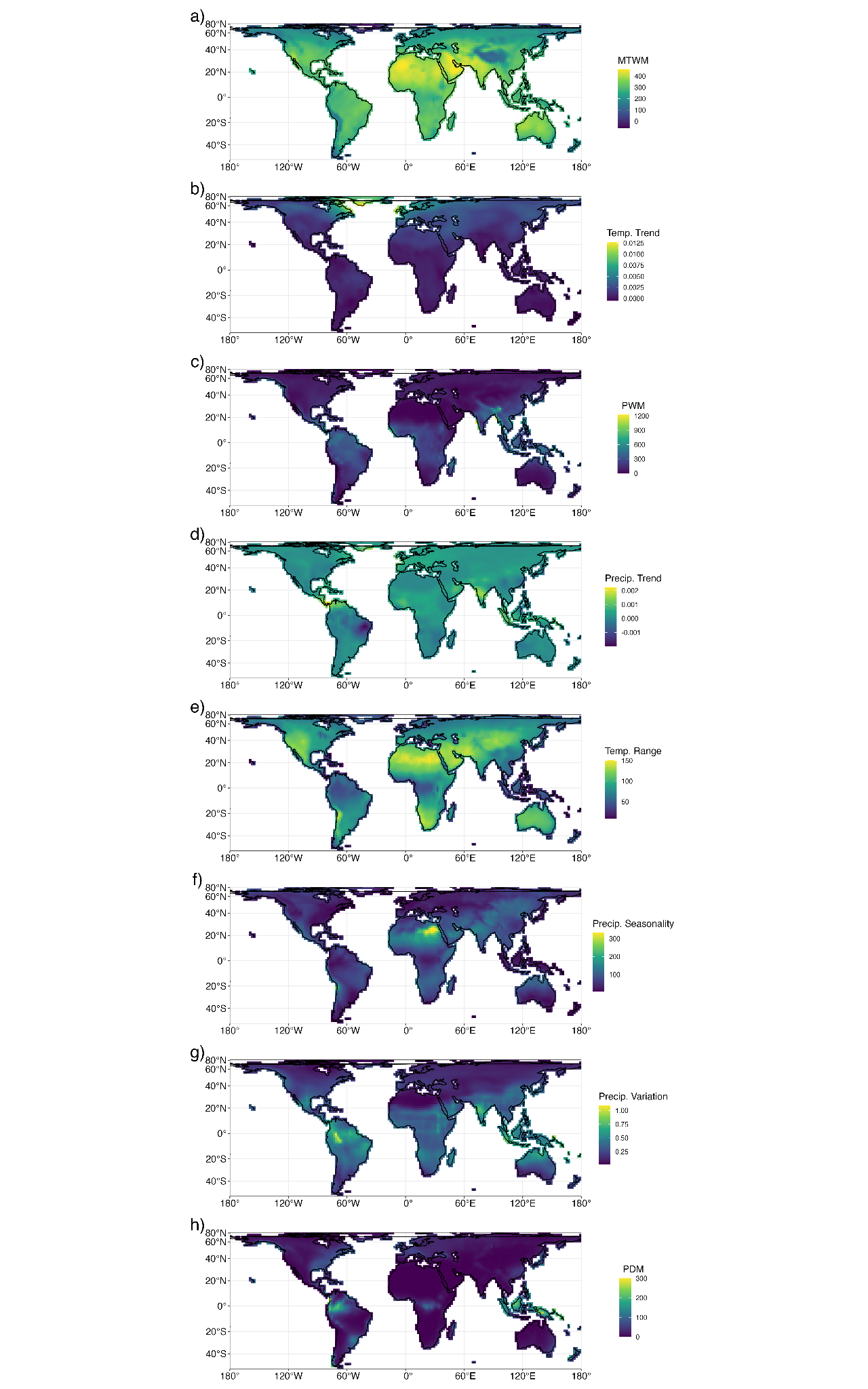

##### Supplementary Fig. 6

The residuals of GDM (a) and GDE (b) for the reported dataset with a minimum OTU threshold of 100. The residuals had no spatial autocorrelation (see Table 2).

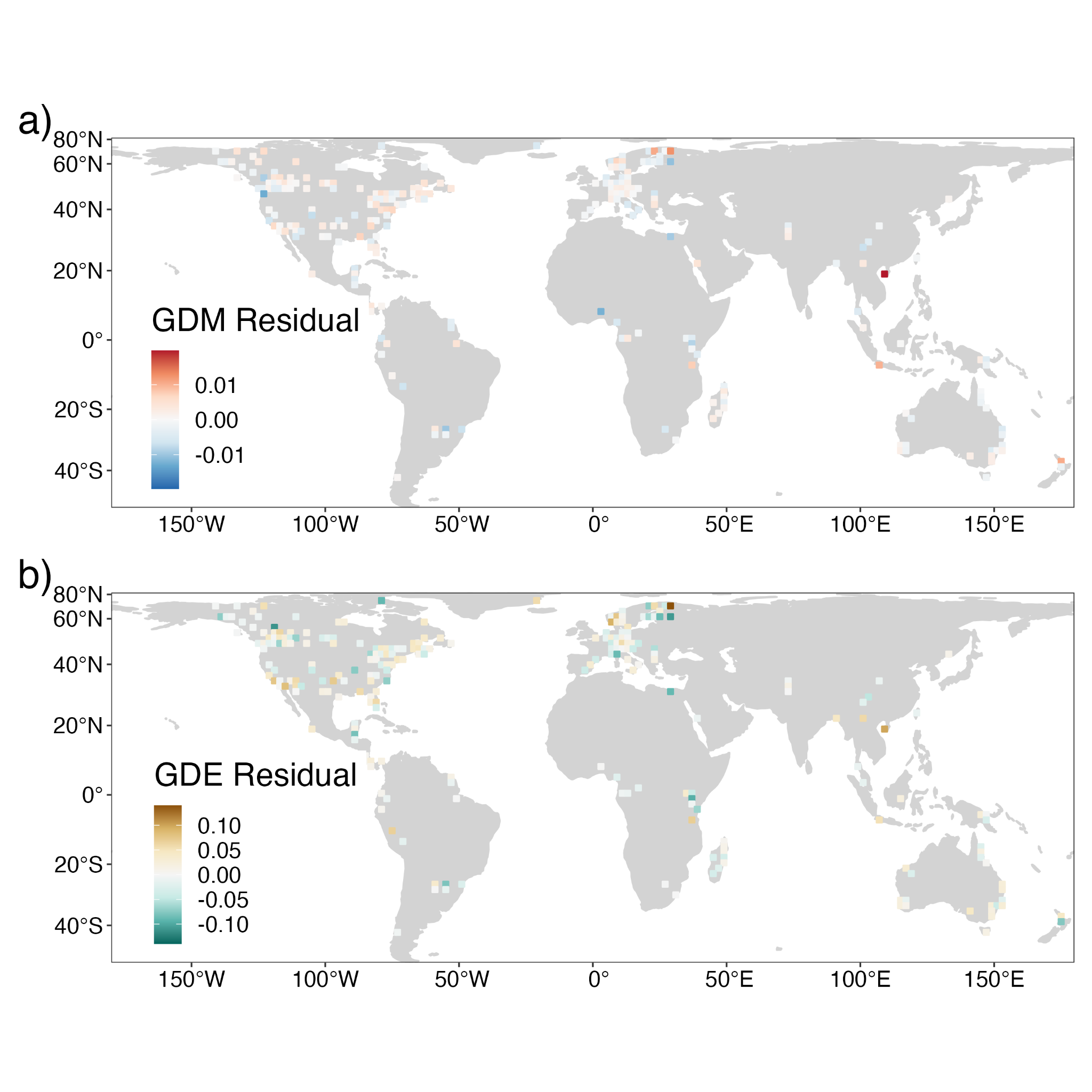

##### Supplementary Fig. 7

Trace plots and prior-posterior overlap plots indicate model stability and parameter identifiability respectively for GDM (a) and GDE (b). The variables considered for GDM (a) are the intercept (B[1]), precipitation trend (B[2]), temperature range (B[3]), maximum temperature of the warmest month (B[4]), precipitation seasonality (B[5]), temperature trend (B[6]), precipitation variation (B[7]), precipitation of the wettest month (B[8]), precipitation of the driest month (B[9]) . The variables considered for GDE (b) are the intercept (B[1]), maximum temperature of the warmest month (B[2]), temperature trend (B[3]), and precipitation of the wettest month (B[4]). Percentages indicate the percentage of the prior distribution that the posterior distribution overlaps with.

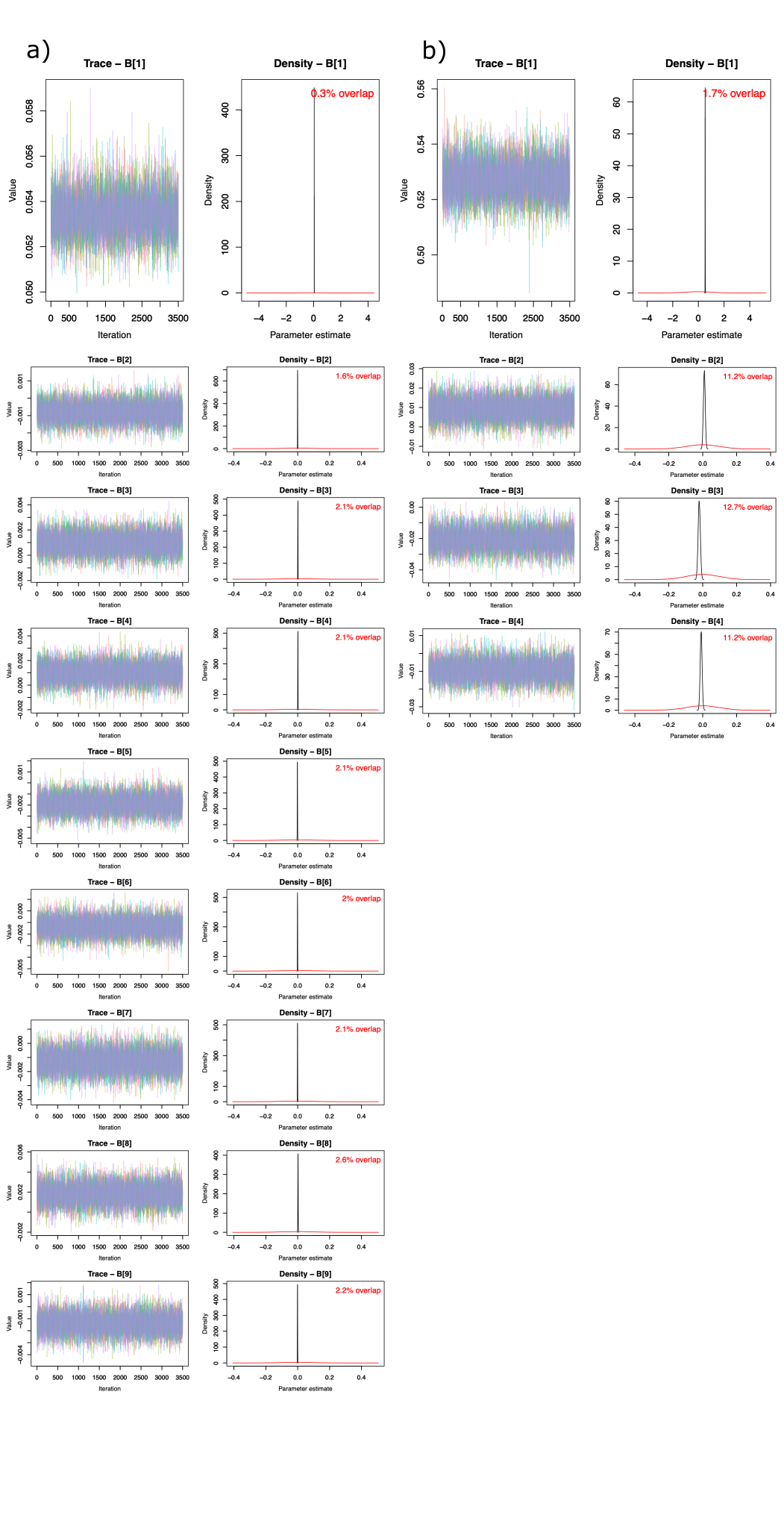

##### Supplementary Fig. 8

Maps showing the global distribution of model uncertainty in GDM (a, b, c) and GDE (d, e, f). The upper 95% highest density interval (HDI) (a, d) and lower 95% HDI (b, e) are plotted. The upper 95% HDI and lower 95% HDI broadly reflect similar patterns to the reported median value (see Fig. 2). We masked in gray areas with environments non-analogous to the environments used for modeling.

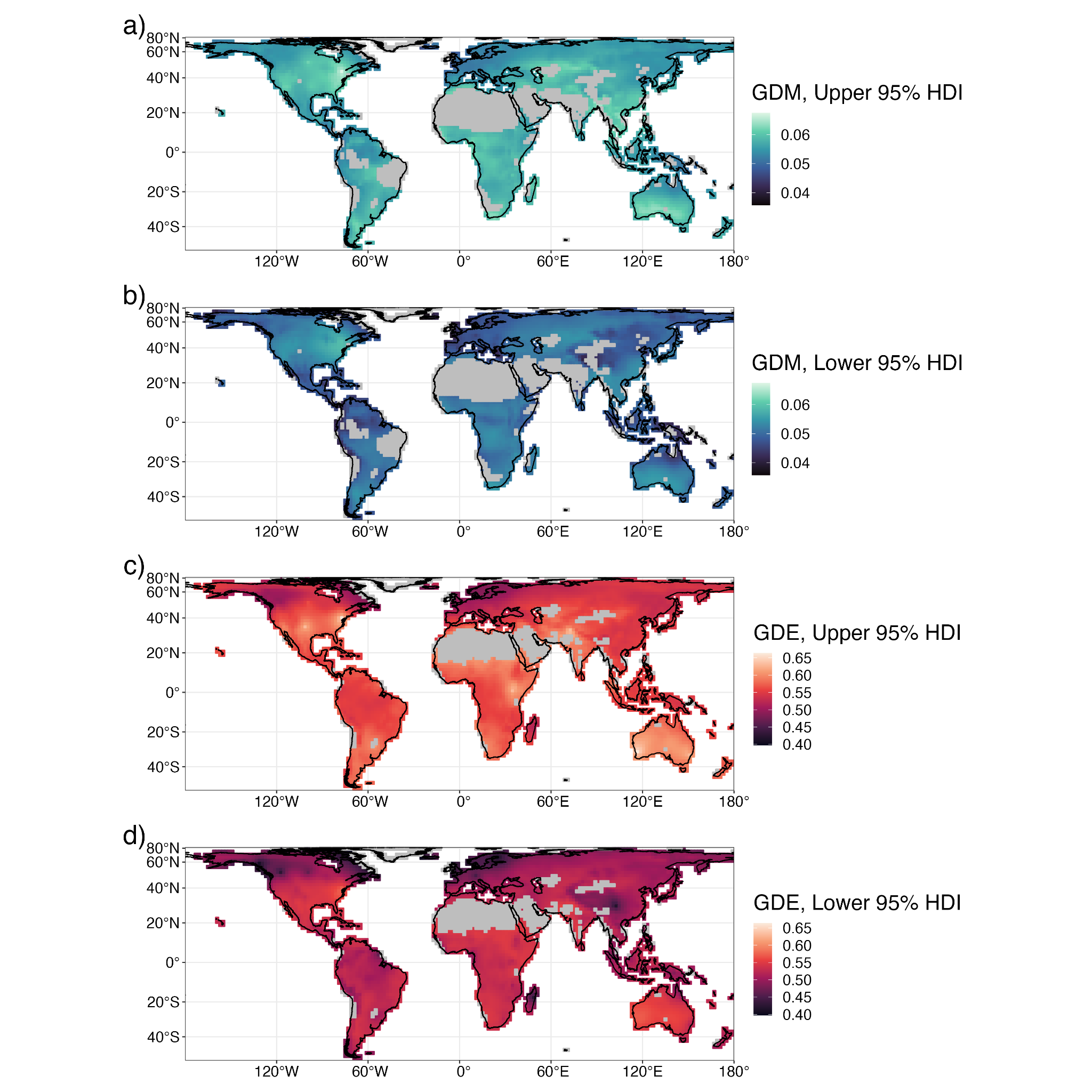

##### Supplementary Fig. 9

Multivariate environmental similarity surfaces, which indicate environmental space that is analogous and non-analogous to that used to train the spatial Bayesian GLMM models of GDE (a) and GDM (b). We masked areas with non-analogous climate in our final projections to limit model extrapolation and uncertainty.

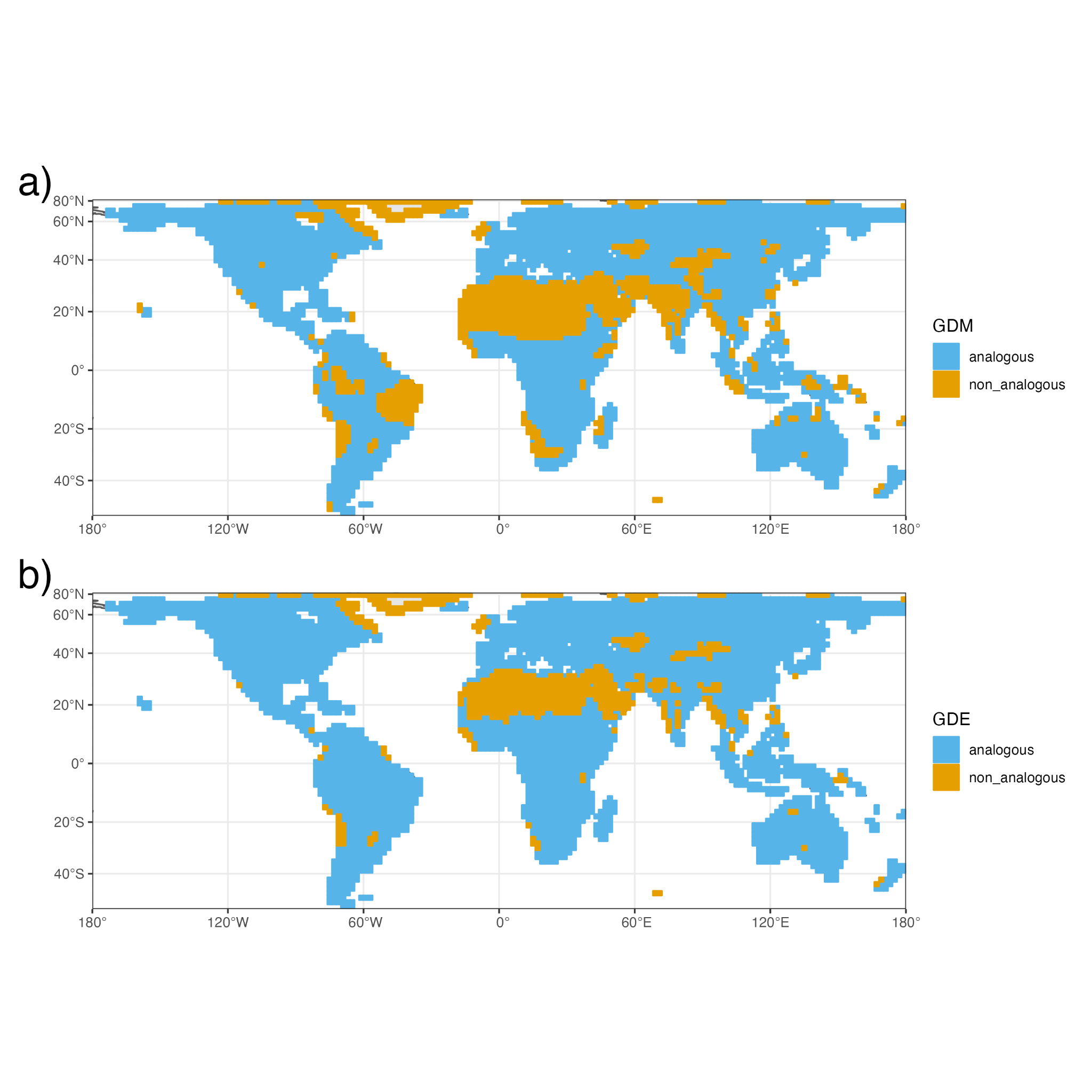

##### Supplementary Fig. 10

The proportion of OTUs belonging to each insect order from the observed data set. Three orders contain over 10% of sampled OTUs each, while 20 orders contain fewer than 5% of sampled OTUs in total (“Other”) (see Supplementary Table 3 for full sampling information).
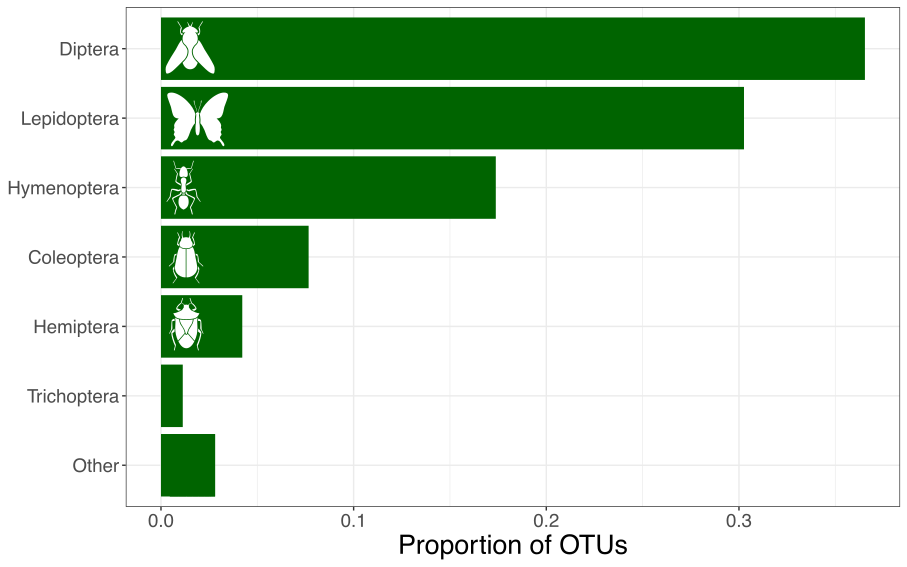

##### Supplementary Fig. 11

The distribution of the number of grid cells each OTU occupies. The tick marks along the x-axis indicate the presence of OTUs that occupy 25 or more cells, but are not visible on the histogram. Over 74 percent of OTUs occupy a single grid cell and over 99 percent occupy fewer than 12 grid cells.

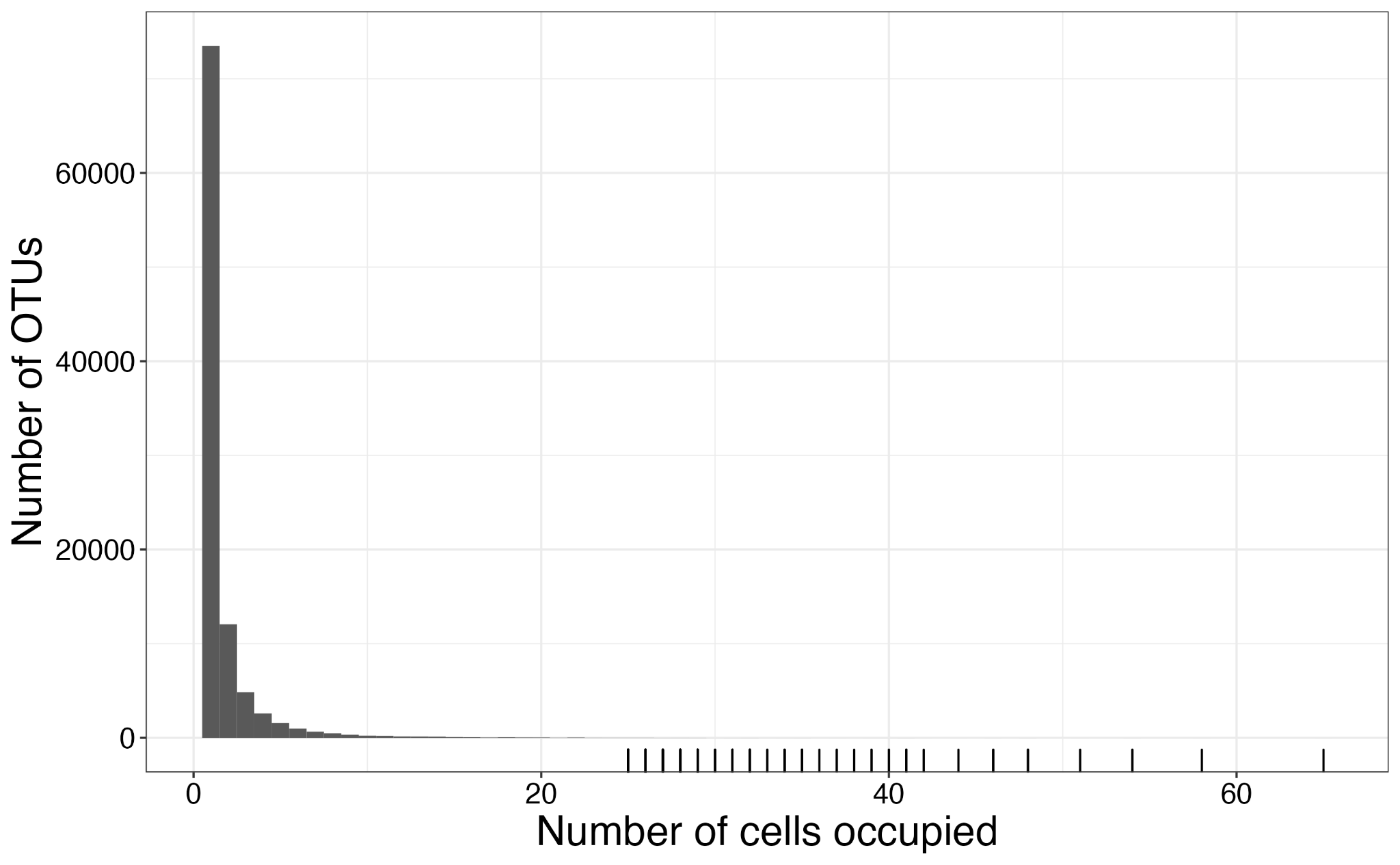

##### Supplementary Fig. 12

The per-cell distributions of GDM (a) and GDE (b) with the most-sampled orders removed compared to the full data set. Three orders included over 10% of sampled OTUs – Diptera (34.0%), Lepidoptera (32.4%), and Hymenoptera (17.3%) (see Supplementary Table 3 for full sampling information). The two datasets with the top three orders removed contained N = 82 cells.

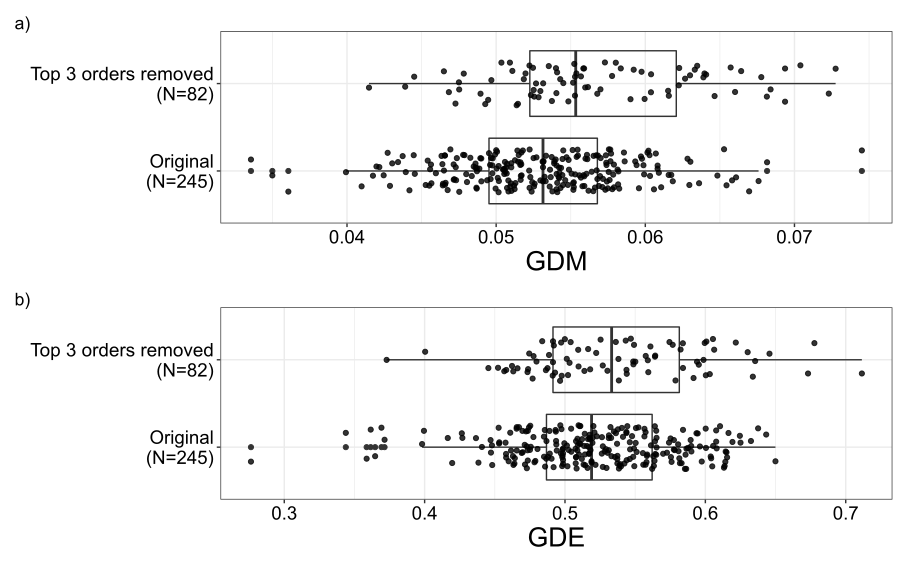

##### Supplementary Fig. 13

The taxonomic distribution of OTU sampling varied geographically. Of the three most sampled orders, Diptera (a) dominated sampling towards the poles and Lepidoptera (b) dominated sampling in the tropics and in some temperate localities at middle latitudes. Hymenoptera (c) typically consisted of fewer than 50% of OTUs sampled per grid cell, although it dominated sampling in Madagascar. Proportions were derived from the number of OTUs sampled per cell belonging to each order.

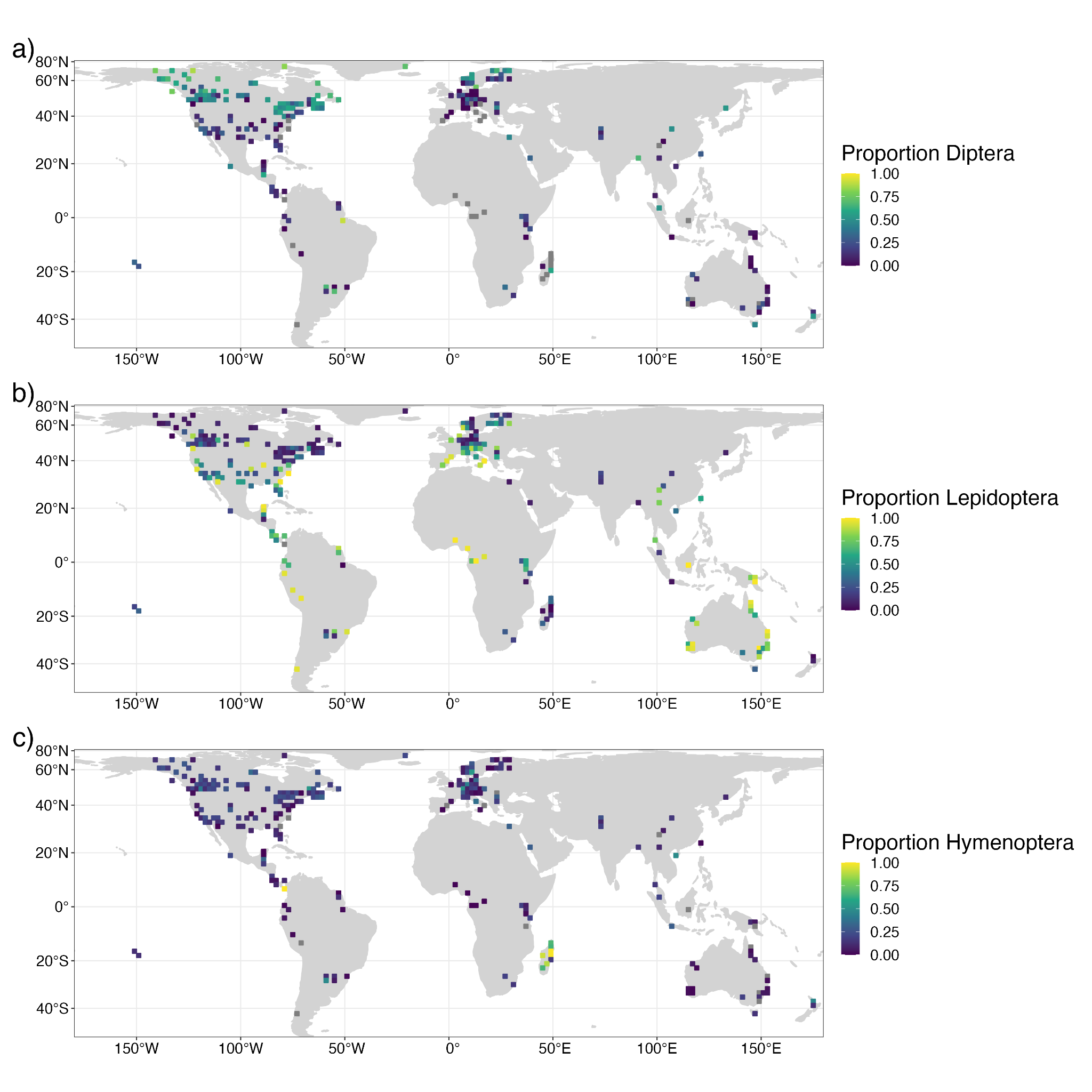

##### Supplementary Fig. 14

The change in the number of grid cells occupied per OTU with latitude. The OTUs are binned, showing the vast majority of OTUs in the data set occupy few grid cells.

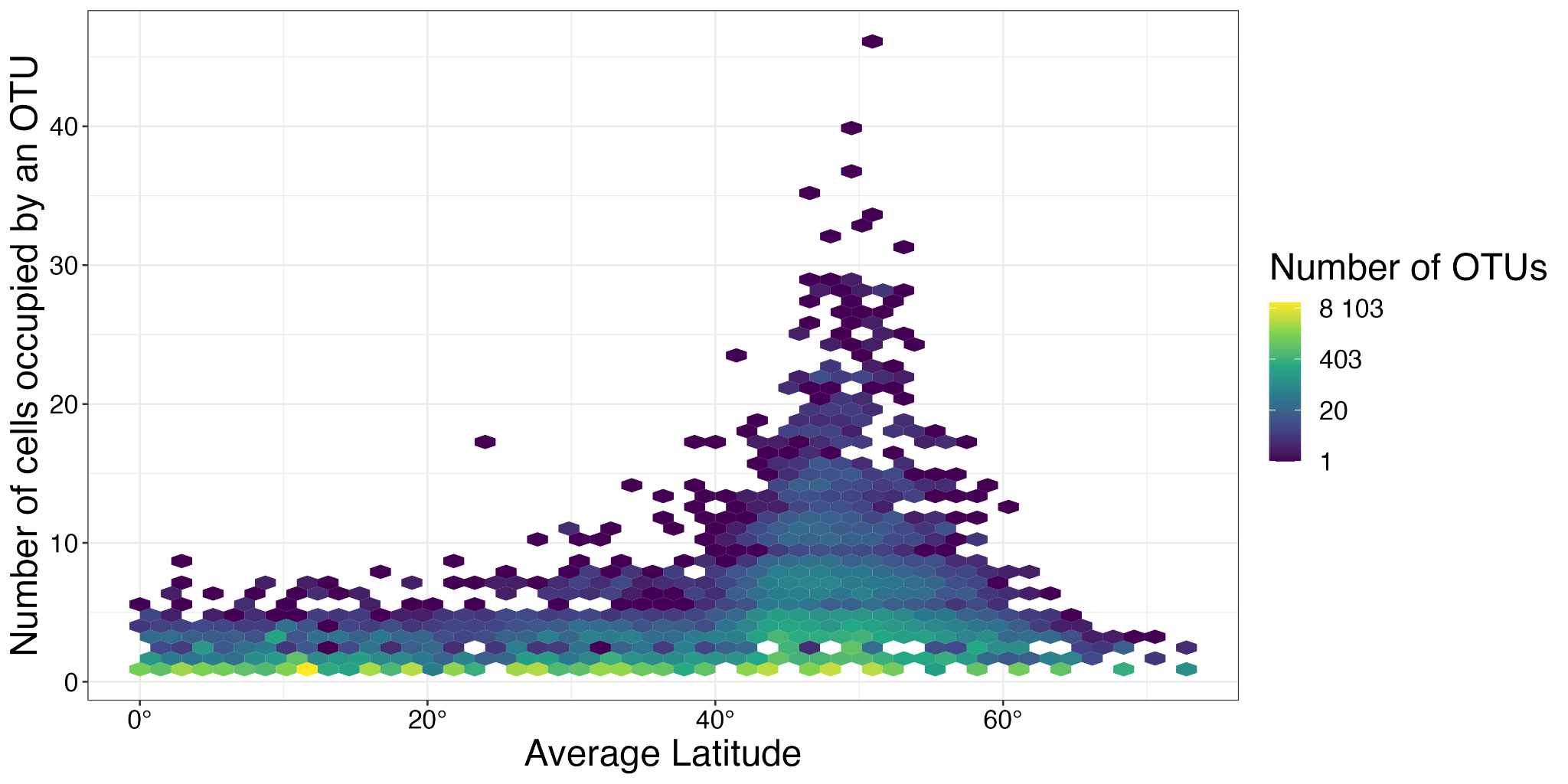

##### Supplementary Fig. 15

Simulations depicting the relationship between nucleotide diversity calculated for all sequences (*pi*) and for only unique haplotypes (*pi**) for 20 values of *Ne*, equally spaced between 1e4 and 1e6. Figures show the average over 1000 simulations of the difference (left panel) or ratio (right panel) between *pi** and *pi* for three sampling regimes representing sample sizes of 5 (blue), 10 (orange), and 50 (green) samples per species. For both smaller *Ne* and larger sample sizes, *pi** is inflated with respect to *pi*, though the proportional size of this difference decreases with increasing *Ne*.
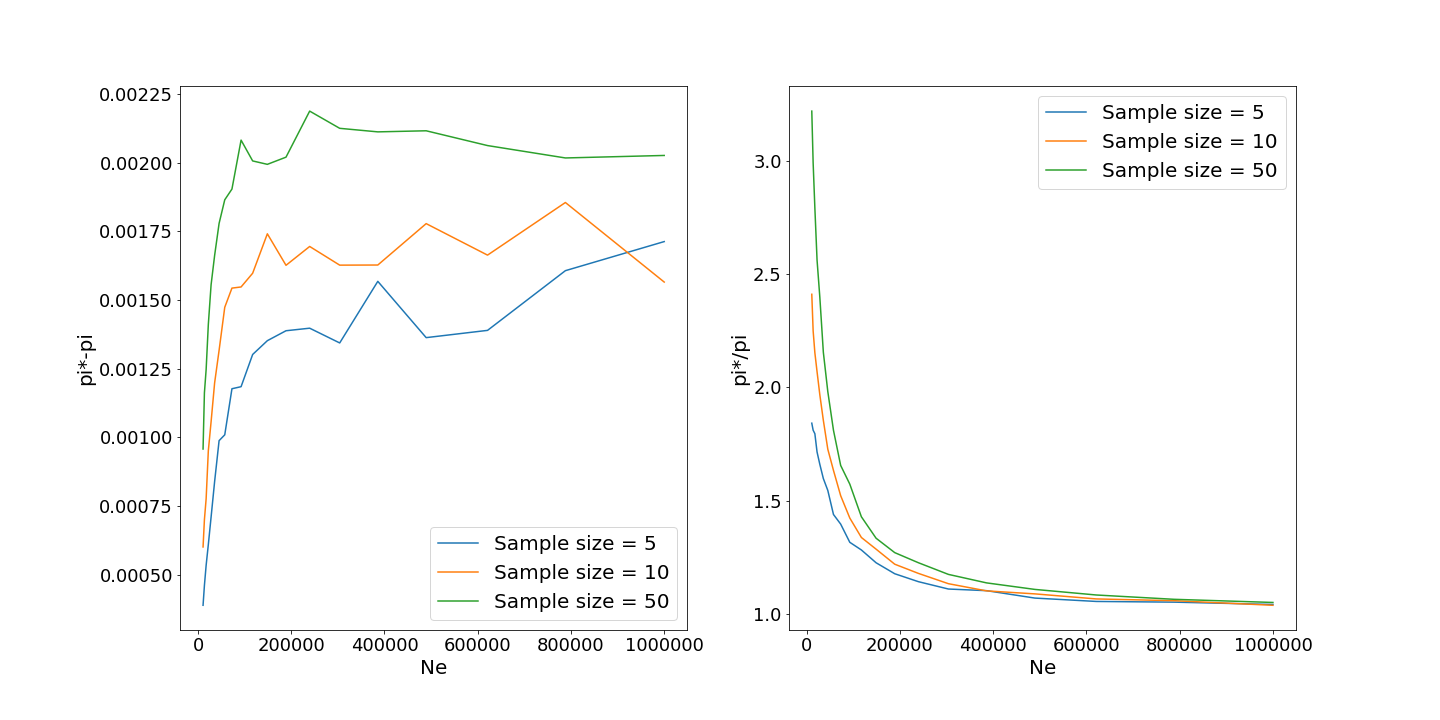

##### Supplementary Fig. 16

Scatterplots of GDM (a,b) and GDE (c,d) and two measures of sampling effort, the number of individuals in a cell (a,c) and the number of OTUs in a cell (b,d) for the minimum OTU threshold of 100. The x-axes are log10-scaled for visualization purposes. No significant correlation exists between the sampling measures and either GD metric (see Supplementary Table 4 ).

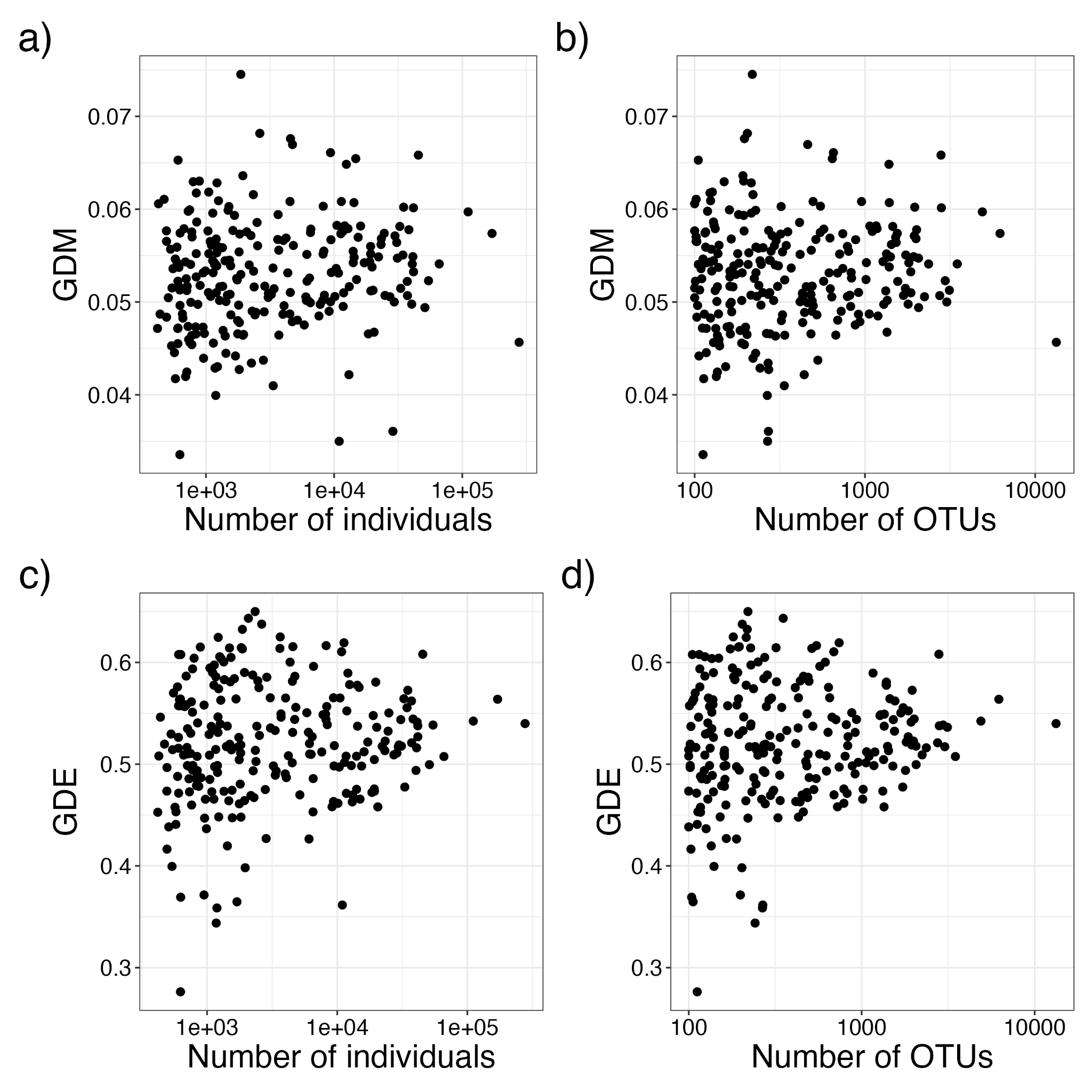

##### Supplementary Fig. 17

Sampling variability in estimates of GDM (a) and GDE (b). We estimated potential sampling variability by resampling the ten assemblages with the largest number of OTUs (2,748 to 13,300 OTUs per grid cell) 1000 times without replacement, with 100 OTUs per sample. The vertical black lines indicate the GD value calculated with the full sample size.

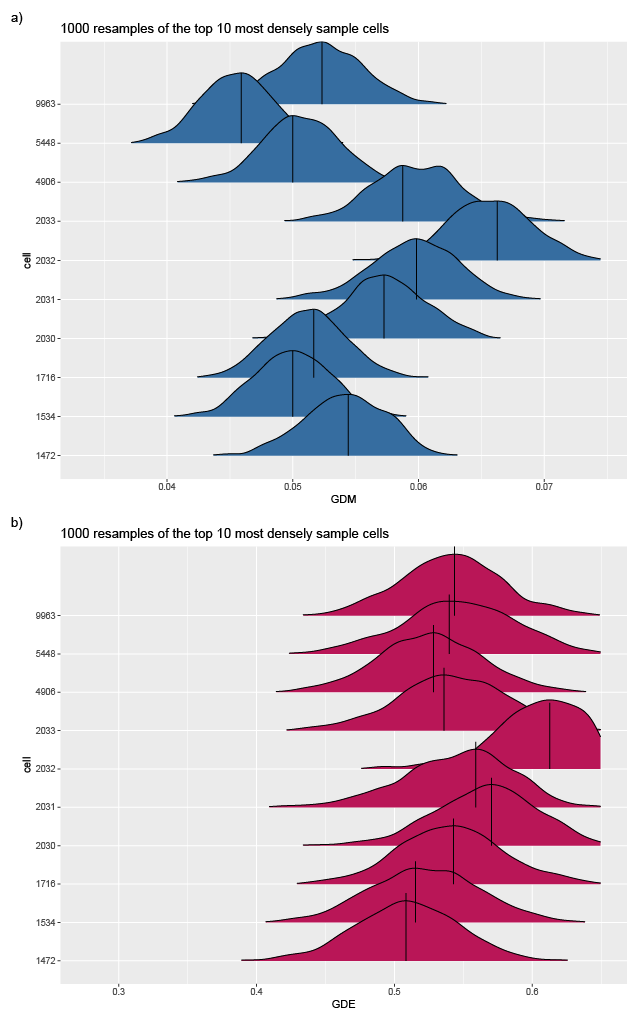

##### Supplementary Table 1

Sampling details for all predictor variables considered for modeling.

| **Variable set** | **Resolution** | **Source** | **Number of variables** | **Time period** | **Citation** |
| --- | --- | --- | --- | --- | --- |
| CHELSA Bioclims | 10 arcmin | <http://www.paleoclim.org/> | 19 | 1981-2010 | ^1^ |
| Global Habitat Heterogeneity | 12.5 arcmin | <https://www.earthenv.org/texture> | 14 | 2001-2005 | ^2^ |
| Elevation | 2.5 arcmin | <https://www.worldclim.org/data/worldclim21.html> | 2 | NA | ^3^ |
| Slope | 2.5 arcmin | <https://www.worldclim.org/data/worldclim21.html> | 4 | NA | ^3^ |
| Dynamic Habitat Indices | 30 arcsecond | <http://silvis.forest.wisc.edu/data/dhis/> | 3 | 2003-2014 | ^4^ |
| Global Human Modification | 30 arcsecond | <https://figshare.com/articles/dataset/Global_Human_Modification/7283087> | 1 | 2016* | ^5^ |
| Climate stability | 1 degree | <https://github.com/spyrostheodoridis/Genetic-geography-of-terrestrial-mammals/blob/master/data.zip> | 4 | 21,000 BP - 1850 AD | ^6^ |

*The time period for the Global Human Modification variable is reported as the median year used by the authors to derive the metric.

##### Supplementary Table 2

Summary statistics from projective prediction feature selection for GDM.

| **Number of predictors** | **Predictors** | **ELPD** | **SE_ELPD_** | **ELPD-DIF** | **SE_ELPD-DIF_** |
| --- | --- | --- | --- | --- | --- |
| 0 |  | 677.8785 | 11.7596 | -10.8082 | 6.0826 |
| 1 | Precipitation trend | 679.4131 | 14.5901 | -9.2736 | 4.7335 |
| 2 | Mean diurnal air temperature range | 678.0864 | 14.2688 | -10.6002 | 3.8210 |
| 3 | Maximum temperature of the warmest month | 680.4610 | 15.0807 | -8.2257 | 3.9907 |
| 4 | Precipitation seasonality | 677.9266 | 15.1074 | -10.7601 | 3.3189 |
| 5 | Temperature trend | 681.6466 | 15.0553 | -7.0400 | 2.9160 |
| 6 | Precipitation variation | 684.5288 | 15.1132 | -4.1579 | 2.7032 |
| 7 | Precipitation of the wettest month | 682.8683 | 14.9238 | -5.8183 | 2.3163 |
| 8 | Precipitation of the driest month | 689.4619 | 14.1291 | 0.7753 | 1.2839 |
| 9 | Temperature variation | 687.9861 | 14.2599 | -0.7005 | 1.0872 |
| 10 | Global habitat heterogeneity (standard deviation) | 687.4668 | 14.3596 | -1.2199 | 1.0756 |
| 11 | Human global habitat modification | 687.2253 | 14.2297 | -1.4614 | 0.8072 |

The ELPD is the expected log (pointwise) predictive density, where a higher ELPD indicates more support for the model. The predictor column is additive, where the model with the predictor listed contains all of the predictors listed before. In addition to ELPD, we report the difference in ELPD (ELPD-DIF), which is the difference in ELPD from the reference model. The SE_ELPD-DIF_ is the standard error of the difference in ELPD from the reference model. SE_ELPD_ is a coarse summary of model support variation that is sensitive to sampling quality, and we take the advice of Vehtari et al. ^7^ to use ELPD-DIF and SE_ELPD-DIF_ when comparing models. We chose the simplest model whose ELPD overlapped with the reference model’s ELPD (ELPD-DIF + SE_ELPD-DIF_ overlaps zero) and corroborated our selection with the heuristic method of selection employed by the *projpred* R package function *suggest_size()*. The chosen model is indicated with gray shading.

##### Supplementary Table 3

Summary statistics from projective prediction feature selection for GDE.

| **Number of variables** | **Predictors** | **ELPD** | **SE_ELPD_** | **ELPD-DIF** | **SE_ELPD-DIF_** |
| --- | --- | --- | --- | --- | --- |
| 0 |  | 253.652 | 11.704 | -28.546 | 8.925 |
| 1 | Maximum temperature of the warmest month | 275.387 | 14.302 | -6.812 | 4.981 |
| 2 | Temperature trend | 277.764 | 15.502 | -4.435 | 4.165 |
| 3 | Precipitation of the wettest month | 279.314 | 17.078 | -2.884 | 3.636 |
| 4 | Temperature variation | 285.854 | 16.165 | 3.655 | 2.258 |
| 5 | Precipitation seasonality | 286.469 | 16.720 | 4.270 | 1.815 |
| 6 | Global habitat heterogeneity (standard deviation) | 284.078 | 17.272 | 1.880 | 1.730 |
| 7 | Precipitation variation | 281.870 | 17.485 | -0.329 | 1.698 |
| 8 | Precipitation trend | 280.279 | 18.175 | -1.920 | 2.218 |
| 9 | Precipitation of the driest month | 280.526 | 17.899 | -1.673 | 1.857 |
| 10 | Mean diurnal air temperature range | 279.379 | 17.846 | -2.820 | 1.767 |
| 11 | Human global habitat modification | 280.342 | 17.869 | -1.857 | 1.780 |

The ELPD is the expected log (pointwise) predictive density, where a higher ELPD indicates more support for the model. The predictor column is additive, where the model with the predictor listed contains all of the predictors listed before. In addition to ELPD, we report the difference in ELPD (ELPD-DIF), which is the difference in ELPD from the reference model. The SE_ELPD-DIF_ is the standard error of the difference in ELPD from the reference model. SE_ELPD_ is a coarse summary of model support variation that is sensitive to sampling quality, and we take the advice of Vehtari et al. ^7^ to use ELPD-DIF and SE_ELPD-DIF_ when comparing models. We chose the simplest model whose ELPD overlapped with the reference model’s ELPD (ELPD-DIF + SE_ELPD-DIF_ overlaps zero) and corroborated our selection with the heuristic method of selection employed by the *projpred* R package function *suggest_size()*. The chosen model is indicated with gray shading.

##### Supplementary Table 4

The raw number and proportion of OTUs belonging to each order in the dataset.

| **Order** | **Number of OTUs** | **Proportion** |
| --- | --- | --- |
| Diptera | 64993 | 0.340 |
| Lepidoptera | 61914 | 0.324 |
| Hymenoptera | 33100 | 0.173 |
| Coleoptera | 14534 | 0.076 |
| Hemiptera | 7918 | 0.041 |
| Trichoptera | 2657 | 0.014 |
| Ephemeroptera | 1020 | 0.005 |
| Orthoptera | 918 | 0.005 |
| Psocodea | 811 | 0.004 |
| Neuroptera | 564 | 0.003 |
| Odonata | 608 | 0.003 |
| Plecoptera | 606 | 0.003 |
| Thysanoptera | 584 | 0.003 |
| Blattodea | 283 | 0.001 |
| Mantodea | 102 | 0.001 |
| Archaeognatha | 27 | 0.000 |
| Dermaptera | 20 | 0.000 |
| Embioptera | 13 | 0.000 |
| Mecoptera | 42 | 0.000 |
| Megaloptera | 61 | 0.000 |
| Phasmatodea | 20 | 0.000 |
| Raphidioptera | 10 | 0.000 |
| Siphonaptera | 33 | 0.000 |
| Strepsiptera | 7 | 0.000 |
| Zygentoma | 1 | 0.000 |
| NA | 46 | 0.000 |

##### Supplementary Table 5

Statistics from Pearson's correlation tests between GD metrics and three summaries of sampling effort.

| **GD Metric** | **Predictor** | ***r*** | ***t*-statistic** | **df** | ***P*-value** |
| --- | --- | --- | --- | --- | --- |
| GDE | Total number of individuals per cell | 0.064 | 1.001 | 243 | 0.318 |
| GDM | Total number of individuals per cell | 0.032 | 0.505 | 243 | 0.614 |
| GDE | Median number of individuals per OTU | 0.060 | 0.944 | 243 | 0.318 |
| GDM | Median number of individuals per OTU | -0.030 | -0.467 | 243 | 0.641 |
| GDE | Number of OTUs per cell | 0.080 | 1.243 | 243 | 0.212 |
| GDM | Number of OTUs per cell | 0.043 | 0.663 | 243 | 0.508 |

### **Supplementary Methods**

### Effect of duplicate alleles on genetic diversity

We used coalescent simulations in *msprime* ^8^ to explore the potential bias in calculation of nucleotide diversity (hereafter *pi*; ^9^) where duplicate alleles may or may not be present in the data. The goal of this simulation study was to replicate data which may be obtained from online databases such as BOLD, where duplicate haplotypes within a focal species may not be represented by duplicate sequence records within the database, potentially biasing estimates of *pi*. For all simulations we set the sequence length to 500bp and the mutation rate to 1e-8/bp/generation, to approximate the features of a typical arthropod COI barcoding dataset. We explored combinations of three different sample sizes (*ss* = 5, 10, 50) to replicate shallow, moderate, and deep sampling of individuals, and a range of effective population sizes (*Ne*) for 20 values of *Ne* equally spaced between 1e4 and 1e6, thus capturing a wide range of *pi* values on the same order as those reported in the literature for arthropods ^10^. We performed 1000 simulation replicates for each combination of sample size and *Ne*, calculated nucleotide diversity for all samples (*pi*) and for only those samples bearing unique haplotypes (*pi**), and recorded the value of the difference between (*pi** - *pi*) and the ratio between (*pi**/*pi*) these quantities averaged across all simulations.

Our coalescent simulation experiments showed a small but consistent upward bias in *pi** with respect to *pi* (Supplementary pi* Fig.). The average absolute difference between *pi** and *pi* increased with increasing *Ne* values uniformly for all sample sizes, while larger sample sizes showed consistently greater difference in *pi**-*pi* across the spectrum of *Ne* values examined. In contrast, the relative difference between *pi** and *pi* decreased with increasing *Ne*, with *pi**/*pi* approaching 1 as *Ne* approached 1e6 for all sample sizes. We draw three main conclusions from these results: 1) The direction of the bias is consistent (*pi** > *pi*) and predictable across a range of *Ne* values; 2) The bias is small for large *Ne* (> 1e6) which is typical for arthropods; 3) For small *Ne* (< 1e6), small sample sizes reduce both the absolute and relative difference between *pi** and *pi*. Taken together, and given the sample sizes and presumed *Ne* values of species from our empirical datasets, we conclude that the bias introduced by sampling only unique haplotypes should in most cases be small and predictable, and that, therefore, it should not unduly influence either the results or our main conclusions.

### Modeling methods

We used projective prediction feature selection (see Methods) for variable selection ^11^. This method projects the posterior information in the full model to smaller submodels by replacing the posterior distribution of the full model with a simpler distribution that is restricted by constraining relevant model parameters. In this case, the regression coefficients decline to zero. The predictive accuracy of these models was evaluated with an approximate leave-one-out (LOO) cross-validation technique, which avoids repeatedly fitting the full model by using an importance weighting scheme outlined in ^7,11^. Under this framework, we used the expected log predictive density (ELPD), specifically the difference in ELPD from the reference model, as the utility function instead of other utility functions (*e.g*., mean squared error) because it measures how well the predictive uncertainties are calibrated, in addition to measuring point predictions. While incorporating spatial autocorrelation into this variable selection step would be ideal, the full 11 variable set contains variables that do not necessarily vary linearly with the response and necessitates an exceedingly complex spatial covariance matrix, which results in the model failing to converge and displaying poor model diagnostics. Therefore, we opted for the simpler non-spatial approach to reduce the parameter space included in the final model.

Rather than estimating random effects at each location, which can be computationally intensive, a spatial field of correlated random effects at a subset of locations or “knots” was modeled. The choice of the number of knots is somewhat subjective, so we fit models using 5, 10, 15, 20, 25, and 30 knots. Models with 20 knots had the lowest residual SAC for GDE, while 30 knots had the lowest residual SAC for GDM, so we chose 20 knots for the GDE models and 30 knots for the GDM models. We used regularizing priors on all slope parameters (N(0, 0.1)) and sigma (N(0, 1)), as well as on the Gaussian process θ parameter (N(0, 5)), which controls how steeply the correlation between knots declines, and the Gaussian process sigma parameter (N(0, 1)), which controls the amplitude of spatial deviations.

### Modeling output

Model output objects for GDM and GDE at all minimum OTU thresholds (10, 25, 50, 100, 150, 200) are available at <https://doi.org/10.6084/m9.figshare.c.6563836.v1>. A guide for loading and interpreting the output is included in the repository and <https://github.com/connor-french/global-insect-macrogenetics>.

### Correlation matrices

Correlation matrices of the proposed 49 variables are at <https://doi.org/10.6084/m9.figshare.c.6563836.v1> as an Excel .xlsx workbook.

### **References**

1.	Karger, D. N. *et al.* Climatologies at high resolution for the earth’s land surface areas. *Scientific Data* **4**, 170122 (2017).

2.	Tuanmu, M.-N. & Jetz, W. A global, remote sensing-based characterization of terrestrial habitat heterogeneity for biodiversity and ecosystem modelling. *Glob. Ecol. Biogeogr.* **24**, 1329–1339 (2015).

3.	Fick, S. E. & Hijmans, R. J. WorldClim 2: new 1‐km spatial resolution climate surfaces for global land areas. *Int. J. Climatol.* **37**, 4302–4315 (2017).

4.	Hobi, M. L. *et al.* A comparison of Dynamic Habitat Indices derived from different MODIS products as predictors of avian species richness. *Remote Sens. Environ.* **195**, 142–152 (2017).

5.	Kennedy, C. M., Oakleaf, J. R., Theobald, D. M., Baruch-Mordo, S. & Kiesecker, J. Managing the middle: A shift in conservation priorities based on the global human modification gradient. *Glob. Chang. Biol.* **25**, 811–826 (2019).

6.	Theodoridis, S. *et al.* Evolutionary history and past climate change shape the distribution of genetic diversity in terrestrial mammals. *Nat. Commun.* **11**, 2557 (2020).

7.	Vehtari, A., Gelman, A. & Gabry, J. Practical Bayesian model evaluation using leave-one-out cross-validation and WAIC. *Stat. Comput.* **27**, 1413–1432 (2017).

8.	Baumdicker, F. *et al.* Efficient ancestry and mutation simulation with msprime 1.0. *Genetics* **220**, (2022).

9.	Nei, M. & Li, W. H. Mathematical model for studying genetic variation in terms of restriction endonucleases. *Proc. Natl. Acad. Sci. U. S. A.* **76**, 5269–5273 (1979).

10.	Leffler, E. M. *et al.* Revisiting an old riddle: what determines genetic diversity levels within species? *PLoS Biol.* **10**, e1001388 (2012).

11.	Piironen, J., Paasiniemi, M. & Vehtari, A. Projective inference in high-dimensional problems: Prediction and feature selection. *EJSS* **14**, 2155–2197 (2020).
